# Angiotensin II regulates the neural expression of subjective fear in humans - precision pharmaco-neuroimaging approach

**DOI:** 10.1101/2022.05.01.490234

**Authors:** Ran Zhang, Weihua Zhao, Ziyu Qi, Ting Xu, Feng Zhou, Benjamin Becker

## Abstract

**Background:** Rodent models and pharmacological neuroimaging studies in humans have been employed to test novel pharmacological agents to reduce fear. However, these strategies are limited with respect to determining process-specific effects on the actual subjective experience of fear which represents the key symptom why patients seek treatment. We here employed a novel precision pharmacological fMRI approach that is based on process-specific neuroaffective signatures to determine effects of the selective angiotensin II type 1 receptor (ATR1) antagonist losartan on the subjective experience of fear.

**Methods:** In a double-blind, placebo-controlled randomized pharmacological fMRI design n = 87 healthy participants were administered 50mg losartan or placebo before they underwent an oddball paradigm which included neutral, novel and fear oddballs. Losartan effects on brain activity and connectivity as well as on process-specific multivariate neural signatures were examined.

**Results:** AT1R blockade selectively reduces the neurofunctional reactivity to fear-inducing visual oddballs in terms of attenuating dorsolateral prefrontal activity and amygdala-ventral anterior cingulate (vACC) communication. Neurofunctional decoding further demonstrates fear-specific effects given that ATR1 blockade (1) reduces the neural expression of subjective fear, but not threat or non-specific negative expressions, and (2) does not affect reactivity to novel oddballs.

**Conclusions:** These results show a specific role of the AT1R in regulating subjective fear experience and demonstrate the feasibility of a precision pharmacological fMRI approach to the affective characterization of novel receptor targets for fear in humans.

## Introduction

Anxiety disorders represent the most common mental disorder worldwide (1) and have become a leading cause of disability (2, 3). Pharmacological treatment approaches – primarily targeting monoamine and GABAergic pathways – have been established, but a considerable number of patients do not adequately respond to the anxiolytic agents or experience negative side effects (4, 5). New treatments targeting alternative pathways to alleviate key symptoms of anxiety disorders are therefore urgently needed.

Anxiety disorders represent a heterogenous group of disorders, with separable key symptomatic and neurobiological alterations (6). Fear-related anxiety disorders (e.g. phobias and social anxiety disorders) and post-traumatic stress disorder (PTSD) are characterized by exaggerated fear responses to potential threats, hypervigilance and avoidance behavior. Animal and humans studies suggest that these symptoms may be rooted in exaggerated threat reactivity or vigilance and an impaired ability to extinguish an acquired fear response (7-9). Translational models that aim at determining new treatments for fear-related disorders therefore commonly test novel compounds based on their efficacy to modulate these domains in rodent models and engage corresponding target circuits in human pharmacological studies with concomitant functional magnetic resonance imaging (pharmaco-fMRI; 10). During the last decade several new target pathways have been explored, including not only classical neurotransmitter systems but also neuropeptides including e.g. the oxytocin and angiotensin-renin system (overview see e.g. 11, 12). Accumulating evidence f suggests that the renin-angiotensin system (13) – classically known for its role in the regulation of blood pressure –represents a promising target to regulate fear (14). Clinical studies have identified an important role of the angiotensin II system in stress- and anxiety-related disorders (15) and population-based studies suggest that pharmacological blockade of the angiotensin II type 1 receptor (ATR1) during trauma exposure decreases the incidence of subsequent post-traumatic fear symptoms (16). Experimental studies utilized the selective competitive ATR1 antagonist losartan to demonstrate that ATR1 blockade facilitates fear extinction in rodents (17) with subsequent pharmaco-fMRI studies in humans demonstrating that losartan has the potential to modulate threat-related amygdala functioning and circuits, including amygdala reactivity to threatening stimuli (13), amygdala-prefrontal connectivity during fear extinction (18) and amygdala-hippocampal connectivity during fear memory formation (19). However, despite the important role of the conditioning paradigm to map fear processing in translational models and despite the important role of the amygdala in fear processing (e.g. 20, 21) it has become increasingly clear that the subjective feeling of fear is not accessible in rodent models and neural indices restricted to isolated brain regions are insufficient to make inferences about mental processing in humans including the subjective experience of fear (22-24). In contrast, the exaggerated subjective feeling of fear is the primary reason for patients to seek treatment, and effects in this domain may represent the most relevant outcome to determine the potential of novel pharmacological treatments for fear (25).

While it remains unclear whether AT1R blockade via losartan (1) modulates fear experience in humans the conventional pharmaco-fMRI approaches are of limited sensitivity to determine process-specific modulations. The commonly applied mass univariate analytic strategies are generally characterized by low spatial and behavioral specificity (see e.g. 24, 26) and ‘classical’ neuroimaging indices of fear reduction such as pharmacologically attenuated amygdala reactivity have been observed during several paradigms including threat conditioning and extinction, aversive anticipation, general negative affect but also various social processes such as face processing or social sharing (e.g. 27, 28-30). Together with studies reporting intact fear and anxiety experience in patients with complete bilateral amygdala damage (20, 31, 32) and studies reporting fear and anxiety decreases in the absence of effects on amygdala activation (e.g. 18, 33, 34), these findings challenge the conventional pharmaco-fmri approach and suggest that behaviorally relevant neurofunctional indices are required to determine precise neurobehavioral profiles of neurotransmitter systems.

Against this background, the present pharmaco-fMRI study aimed to determine the specific neurofunctional effects of LT on fear vigilance by means of capitalizing on process-specific multivariate neural signatures for fear and associated processes. These multivariate signatures, which have been developed to predict emotional processing, can provide larger effect sizes in brain-outcome associations as compared to local multivariate and univariate models (35) and thus represent more precise and comprehensive brain-level descriptions of mental processes (24). Moreover, the multivariate signatures allow to access the individual mental and emotional state directly based on neurofunctional activation patterns without the need for subjective self-report. This has the advantage of avoiding meta-reflective processes which may confound the actual mental process of interest and limit ecological validity. In the current study, we, therefore, employed a number of emotional process-specific multivariate signatures to decode the effects of LT on fear-related emotional experiences, including general negative affect (36), acquired threat (or conditioned fear (37)) and subjective fear experience (23). The combination of these decoders allowed us to test which specific fear-related processes are modulated by the AT1R system. To trigger fear-related neurofunctional processes while controlling for higher-order cognitive evaluation processes we employed a visual oddball paradigm including fearful oddballs to trigger fear reactivity and vigilance (38). The paradigm additionally included non-fearful novel oddballs and target oddballs to determine non-specific treatment effects on novelty responses and general attention.

Based on accumulating evidence from animal and human studies suggesting a role of the AT1R system in regulating fear- and threat-related processes (13, 18, 39-41), we hypothesized that LT would reduce neural vigilance towards fearful oddballs as reflected by attenuated neural reactivity and connectivity in the fronto-limbic fear circuitry. Based on recent evidence suggesting that ATR1 blockade specifically modulates processing of threat-related – but not neutral or salient positive – stimuli (18, 19, 42), we expected that the neurofunctional decoding analyses would reveal specific effect of LT on the neural expression of subjective fear but not associated affective processes or neural reactivity towards novel oddballs.

## Method and materials

### Participants

Ninety healthy right-handed participants (female=34; mean age=21.17 ±2.12) were enrolled. They were free of current and past mental disorders and medication. Participants were asked to abstain from caffeine and alcohol in the 24h before the experiment. Smokers and subjects with MRI or treatment contraindications were excluded. Due to excessive movement and low performance 3 subjects were excluded, leading to 44 LT and 43 PLC subjects for the final analyses. Participants provided written informed consent, the study had full ethical approval by the local ethics committee and was in line with the Declaration of Helsinki.

### Procedures

The current study employed a randomized placebo-controlled, double-blind, between-subject pharmaco-fMRI design. Participants were randomly assigned to either a single dose (50 mg) of LT or placebo (PLC) p.o. packed in identical capsules. An independent researcher packed and dispensed the capsules according to a computer-generated randomization list (n=2 groups). The paradigm started 90 minutes after treatment administration in line with the peak plasma pharmacodynamics of the drug (43, 44). To control for pre-treatment differences and non-specific effects of treatment participants completed the State-Trait Anxiety Inventory (STAI), Positive and Negative Affect Schedule (PANAS) and assessments of heart rate and blood pressure at baseline, peak plasma and after the experiment (see **Figure 1A**).

**Figure 1.**
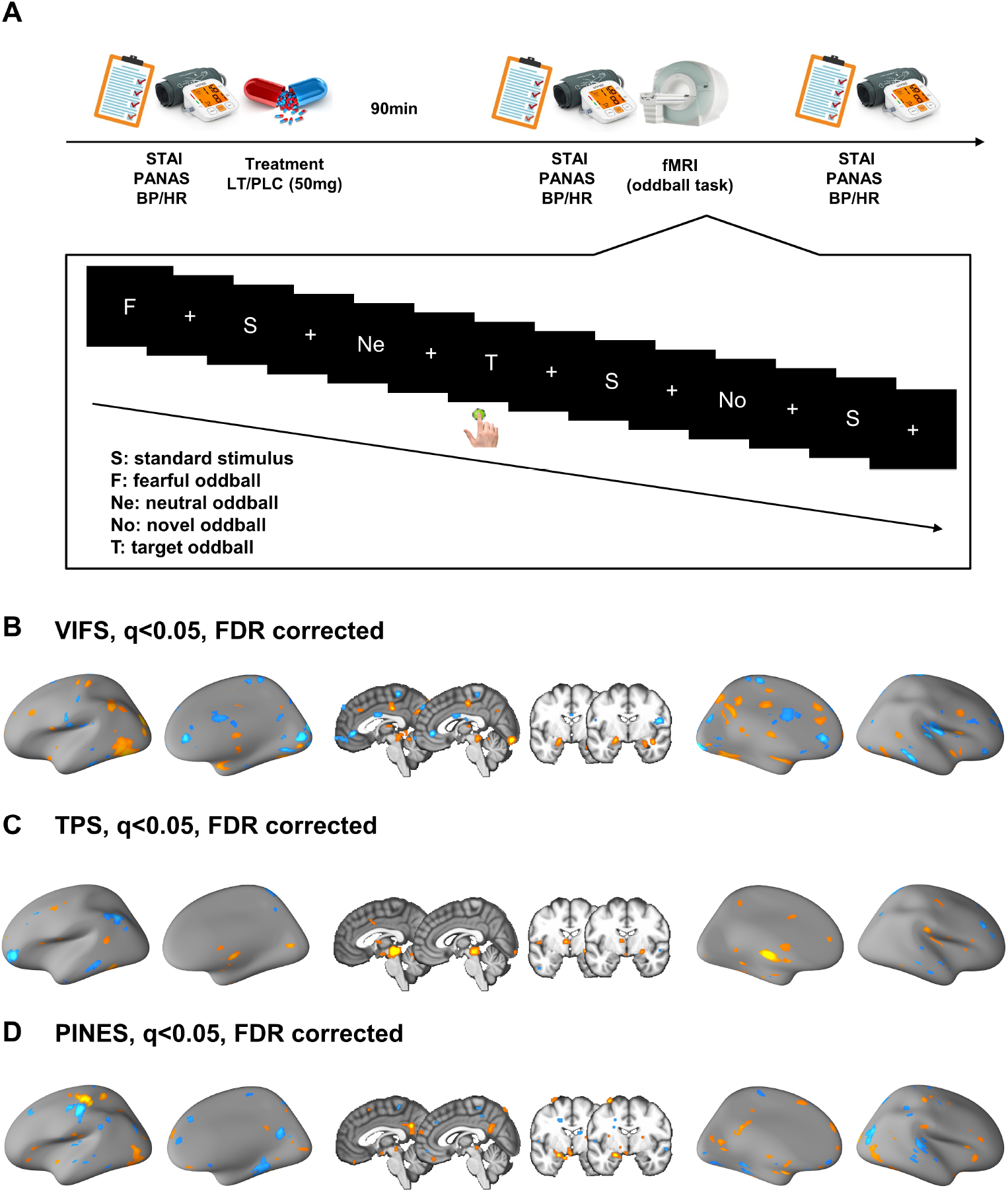
Experimental design and neural decoding of fear-related process. (19) Experimental timeline and schematic synopsis of the fMRI task (oddball paradigm); (19) Visually induced fear signature (VIFS); (19) Threat (conditioned fear) predictive signature (TPS); (D) Picture-induced negative emotion signature (PINES). Displayed at q < 0.001, FDR corrected, for visual purpose.

### Stimuli and oddball paradigm

A modified version of a previously validated oddball fMRI paradigm with the fearful, neutral, target, and novel oddballs was employed (38). Participants were presented with a series of visual stimuli and 66.6% of the stimuli were neutral ‘standard’ images. ‘Oddballs’ occurred with a probability of 8.3% per oddball condition. Neutral and target oddballs employed single neutral animal pictures, the fearful oddball employed a fearful animal picture and the novel oddball used 10 neutral animal pictures with each presented once in a run. Participants completed 3 runs with 120 trials each, resulting in a total of 360 trials (240 standard trials, and 30 oddball trials for each of target, neutral, fearful, and novel stimuli). Each trial consisted of a 500 ms presentation of the picture followed by a white fixation cross on a black background with a mean interstimulus interval (ISI) of 2.7s. ISI was jittered with ±300 ms. The order of stimuli was pseudo-randomly distributed throughout the task. Before the experiment, participants were asked to press a single button with the right index finger only when they saw the ‘target oddball’ during fMRI.

The stimuli for the oddball paradigm were selected from the Nencki Affective Picture System (NAPS) (45) which is a standardized set of 1356 realistic, high-quality photographs including people, faces, animals, objects, and landscapes. For the present paradigm, 34 pictures displaying animals were selected based on ratings in an independent sample (n=60) rating the fear degree of the stimuli on a 1-5 Likert scale. We selected a stimulus with moderately higher fear and arousal induction (depicting a ferocious wolf) than the other stimuli as a fear stimulus to avoid ceiling effects. For a detailed description of the stimuli and ratings see **Supplemental Table S1**. Stimuli were presented using E-Prime stimulus presentation software (Psychology Software Tools, Sharpsburg, PA) a presented via a mirror attached to the head coil enabling viewing of projected images onto a screen positioned at the back of the MRI bore. The paradigm had a total duration of 20 minutes.

### Functional MRI acquisition and preprocessing

Functional MRI data were collected using standard sequences on a 3.0-Tesla MRI system and preprocessing was conducted using validated protocols in Statistical Parametric Mapping (SPM12 v7487, https://www.fil.ion.ucl.ac.uk/spm/software/spm12/)

The first-level general linear model (GLM) included five task regressors. For those participants who missed the target and/or pressed the button during any other conditions, we included additional task regressors to model missing and/or false alarm trials (details see **Supplemental Material**).

### Exploring the treatment effects on brain activity

Based on previous studies showing effects of AT1R blockade on neural responses to fear- and threat-associated stimuli (e.g. Reinecke et al., 2019; Zhou et al., 2019) we hypothesized an effect of LT on vigilance towards fearful stimuli and employed a mass-univariate analysis to test whether the administration of LT had effects on brain activation to fearful versus neutral oddballs. Specifically, we employed a partitioned error ANOVA approach that models the within-subject factor fear vigilance on the first level (fearful oddball >neutral oddball) and the between-subject factor on the second level (treatment, LT vs PLC). This approach may facilitate a more robust modelling in SPM as compared to the pooled error approach (46) and additionally allowed us to align the contrast comparions between the mass-univariate and multivariate decoding analysis focusing on fearful oddball vs neutral oddball. Based on the fronto-insular-limbic networks described in previous studies employing oddball paradigms (38) as well as previous studies with LT reporting effects on amygdala-frontal circuits (13, 18) the analyses focused on these regions as defined by the Harvard-Oxford cortical and subcortical structural atlas (thresholded at 25% probability). To this end, a single mask encompassing the anterior cingulate cortex (ACC), amygdala (Amyg), insula, middle frontal gyrus (referred to as dorsolateral prefrontal cortex (dlPFC)), inferior frontal gyrus (referred to as ventrolateral prefrontal cortex (vlPFC)) and frontal medial cortex (referred to as ventromedial prefrontal cortex (vmPFC)) (see also supplementary **Figure 1** for the mask) was used for the correction of multiple comparisons, with a threshold of threshold-free cluster enhancement (TFCE) *p* <0.05, family-wise error (FWE)-corrected at peak-level.

We found LT effects on fearful, but not neutral, oddballs (see Results). However, treatment-induced changes in brain activity do not necessarily indicate that the treatment affected the subjective fear experience. For example, the fearful oddball stimulus can also induce other emotional experiences (e.g., salience, arousal and negative valence) which are inherently associated with fear. The observed effect may be therefore due to non-specific treatment effects on basal processes of salience or emotional experience. We, therefore, asked whether LT has a specific effect on the fear experience. To address this question we (1) tested whether LT had effects on brain activity to the novel oddball which could induce higher salience, but not fear, as compared to neutral oddball, and (2) applied three multivariate predictive models, which measured subjective fear, conditioned threat, and general negative emotion separately, to the contrasts of fearful and neutral oddballs. Specifically, we first used the visually induced fear signature (VIFS) (**Figure 1B**) to predict fear experience in the different treatment groups. The VIFS has a robust generalization and high sensitivity to predict fear experience across populations, fear induction paradigms and MRI systems (23). Inference on model performance was performed using permutation testing with 10,000 random shuffles. To test the specificity of the effects we additionally examined effects on the responses of neurofunctional signatures for related processes, in particular non-specific negative affect and threat (or conditioned fear, which is traditionally also referred to as fear). We additionally employed a picture-induced negative emotion signature (PINES) (36) designed to capture a neural activation pattern distributed across the brain that accurately predicts the level of current negative affect induced by visual stimuli, and the threat predictive signature (TPS) (37) which was developed to accurately predict neural threat reactivity to test LT effects on conditioned threat, and general negative emotion separately (**Figure 1C and D**). Notably, all three decoders encompassed the amygdala as a reliably predictive region (see also 23, 36, 37) although emotional prediction based on the whole-brain patterns performed superior to the amygdala restricted predictions.

### Exploring the treatment effects on brain connectivity

Based our findings showing specific effects of LT on fear vigilance, and in the context of previous studies demonstrating the important role of amygdala-prefrontal circuits in anxiety and early fear detection processes (e.g. 20, 21, 31) as well as in mediating some fear and negative information processing related effects of LT (13, 19, 33) we next examined effects of treatment on amygdala functional connectivity during fear vigilance. To this end, we employed a task-based functional connectivity analysis (generalized form of context-dependent psychophysiological interactions; gPPI (47)) with identical first-level contrasts as the activation analyses and the bilateral amygdala from the Harvard-Oxford subcortical structural atlas as the seed region. In this exploratory analysis we specifically focused on the contrast of [fearful oddball > standard] using a voxel-wise comparison at the group level. In line with the activation analysis, correction of multiple comparisons was restricted to the predefined mask (excluding the amygdala which served as the seed region).

### Statistical analysis of behavioral data

Statistical analyses of behavioral data were carried out using SPSS software (version 22; IBM Corp., Armonk, NY). Potential drug-induced changes in mood and physiological parameters were assessed using independent samples *t* tests and Bayesian *t* tests on indices collected during peak plasma. Effects on false alarms and accuracy for the target oddball were analyzed using independent samples *t* tests. All reported *p* values were two-tailed, and *p* < 0.05 was considered significant.

## Results

### Participants

The final sample included n=87 (mean age=21.17 ±2.15) subjects and the treatment groups were well matched on sociodemographic parameters (LT, n=44, 17 females; PLC, n=43, 16 females). There were no effects of LT on non-specific emotional indices or cardiovascular activity (**Table 1**).

**Table 1.**
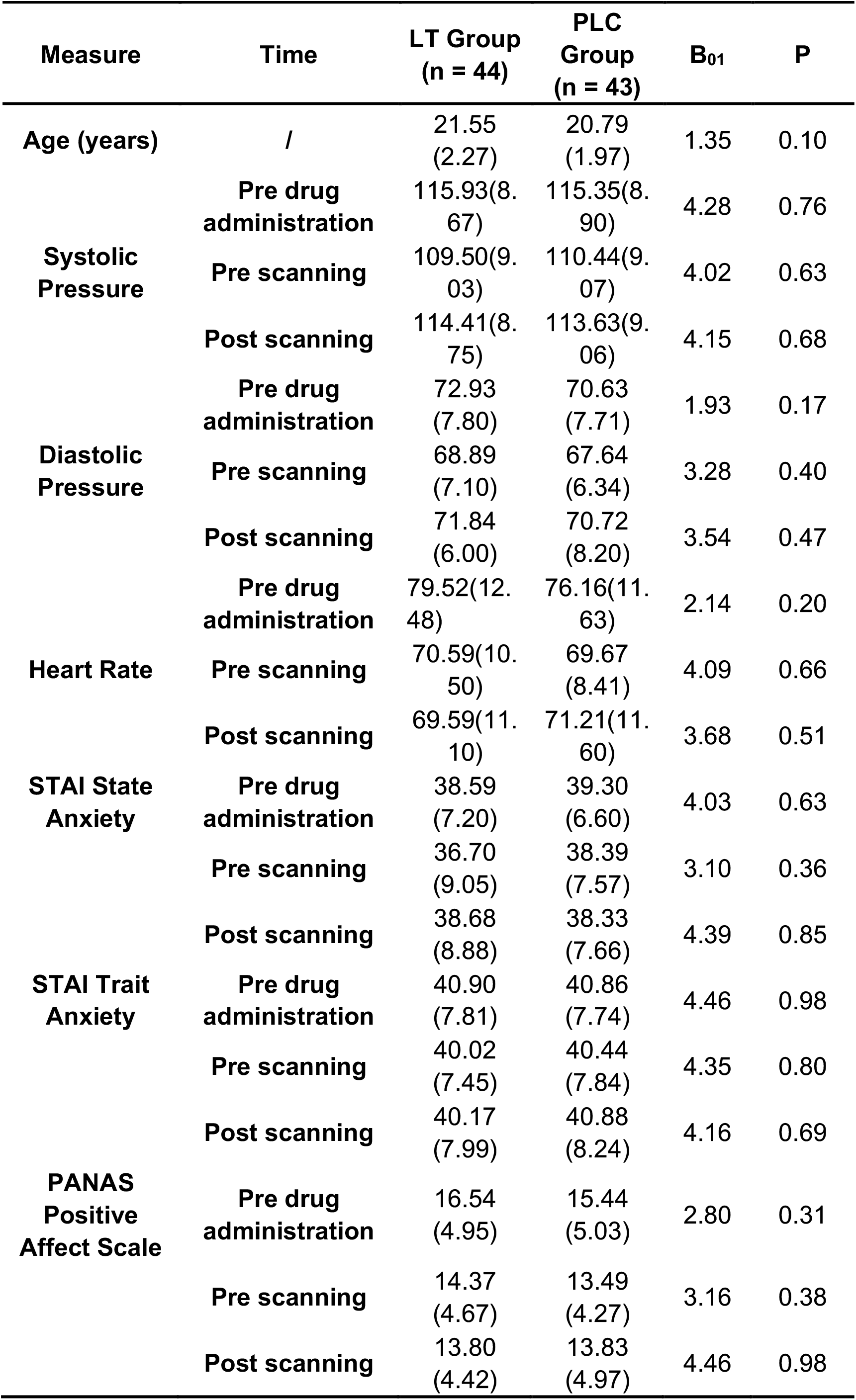
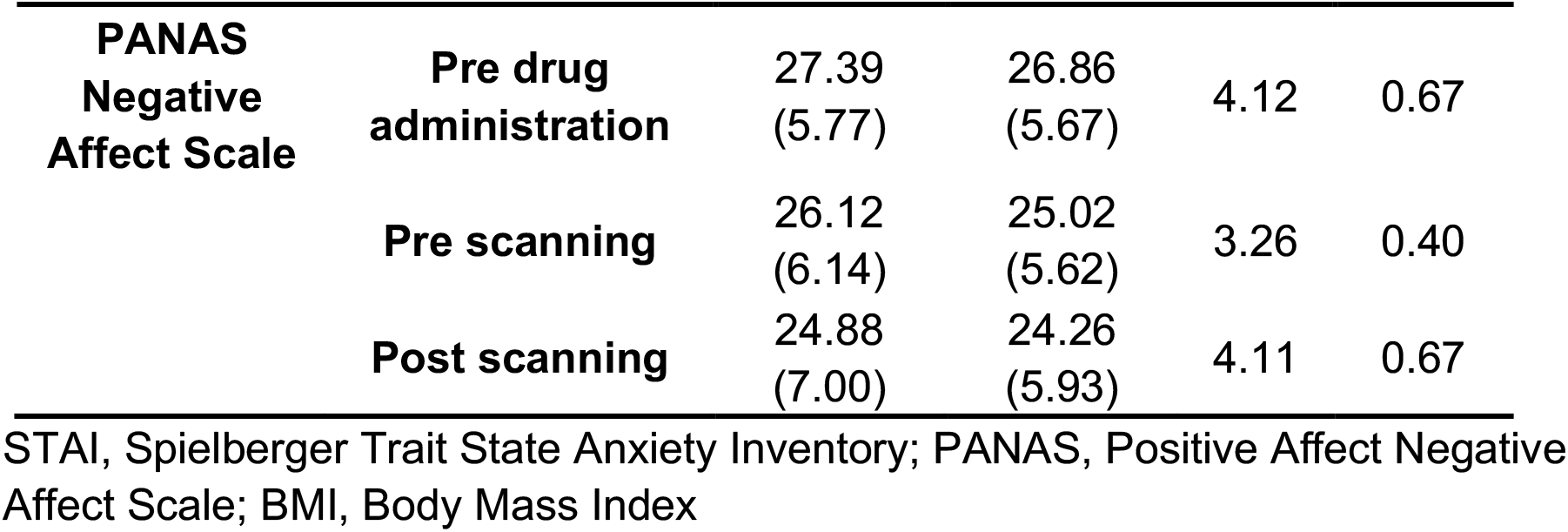
Participant Demographics and Control Measures

### Behavioral indices of attention and task engagement

In order to maintain attentional engagement during the oddball task, participants were required to press a button in response to the target picture. Analyzing the number of false alarms and accuracy rate revealed that both groups showed a high task engagement and accuracy (94% or 95% accuracy, respectively), while LT did not affect this measure of basal attention (all between-group comparisons p >0.1, details provided in **Table 2**).

**Table 2.**
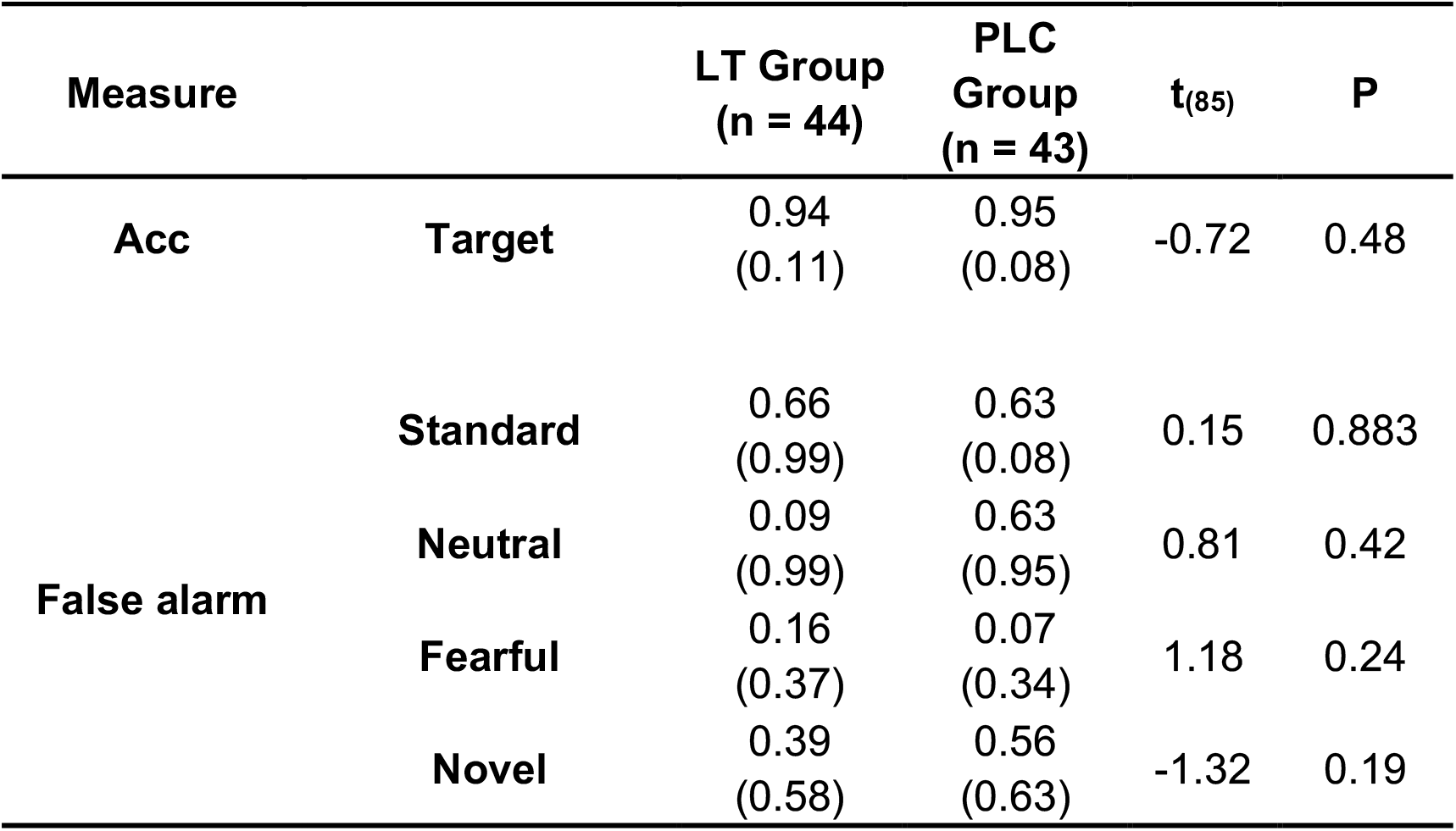
Accurancy and False alarm in Oddball Task

### Effects of AT1R blockade on neural fear reactivity

We initially examined neural engagement during the task irrespective of treatment using whole-brain one-sample t-tests in the entire sample. Results showed significant activation of the amygdala across all oddball contrasts (fearful oddball, neutral oddball novel oddball, and target oddball > standard; TFCE p <0.05, FWE-corrected, displayed in supplementary **Figure 1**). These findings indicate that the amygdala is engaged during rare salient stimuli suggesting an engagement in the detection of salient or novel - and potentially threatening - stimuli in the environment (22).

We next tested whether LT had effects on fear vigilance, and found a significant treatment (LT vs. PLC) and condition (fearful oddball vs. neutral oddball) interaction effect in the left dorsolateral prefrontal cortex (dlPFC; peak MNI x, y, z=-48, 26, 32; t_85_=-3.58; TFCE p=0.04, FWE-corrected at peak level; k=71, **Figure 2A**). Post-hoc two-sample ttests with voxel-wise comparisons within pre-defined mask demonstrated decreased brain activity to fearful, but not neutral, oddballs following LT administration in the left dlPFC (cluster 1: peak MNI x, y, z=-44, 20, 28; t_85_=-3.72; TFCE p=0.02, k=217; cluster 2: peak MNI x, y, z=-30, 16, 44; t_85_=-3.30; TFCE p=0.04, k=53; FWE-corrected at peak level, **Figure 2B**).

**Figure 2.**
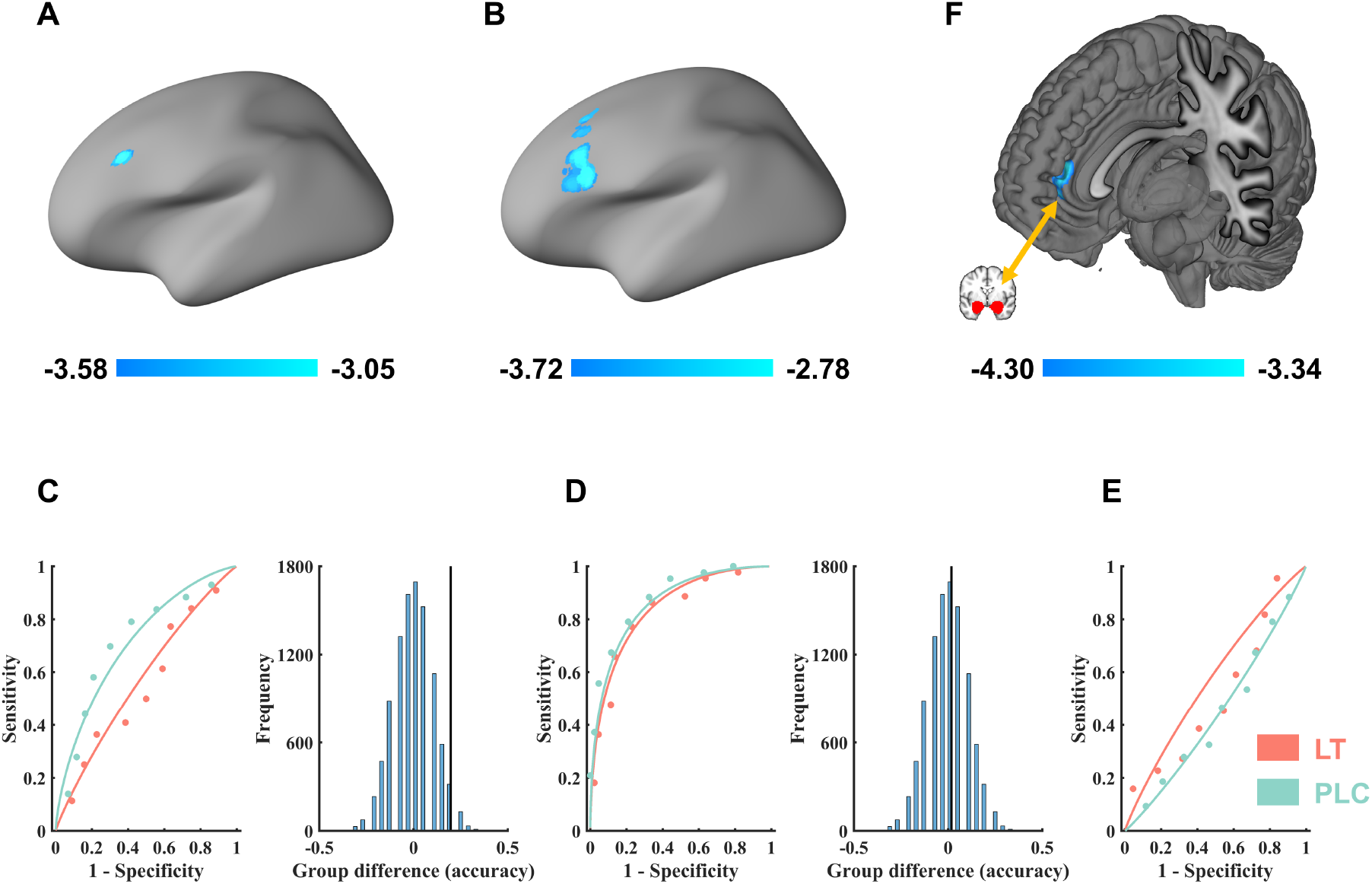
Main results (19) left dorsolateral prefrontal cortex (dlPFC) showed significant treatment (placebo vs. losartan) × condition (fear vs. neutral oddballs) interaction effect; (19) Losartan selectively reduced dlPFC reactivity to the fearful oddball (p < 0.05, TFCE corrected); (19) VIFS discriminated fearful from neutral oddballs in the PLC but not the LT group and classification accuracies differed significantly between the groups (permutation test p < 0.05); (D) PINES discriminated fearful vs. neutral oddballs in both groups with similar accuracies. (19) TPS could not discriminate fearful vs. neutral oddballs in either group; (19) Losartan specifically decreased functional connectivity between ACC and amygdala during the fearful oddball presentation;

### Process-specific effects of AT1R blockade on neurofunctional fear experience

We next employed two independent methods to test the specificity of the LT effects on fear experience. We first tested whether LT would also affect basal processes that are inherently associated with fear vigilance such as salience processing. To this end we performed an identical mass-univariate analysis of treatment effects (LT vs. PLC) on novelty (novel vs. neutral oddballs). We hypothesized that if the observed treatment effects were driven by effects on basal salience processing we should also find brain regions exhibiting significant interaction effects. However, no brain region showed significant interaction effects (TFCE p < 0.001, k≥10), which suggest that the effects of LT on fearful oddball are not due to effects on basal salience processing. Moreover, we applied the emotion-specific neurofunctional signatures to predict fearful oddballs from neutral oddballs in the separate treatment groups. Application of the VIFS capturing the level of subjective fear experience revealed significant classification performance in the PLC (accuracy=70 ±7.0%, binomial test p=0.01) but not the LT group (accuracy=50 ±7.5%, binomial test p=1, **Figure 2C**). Moreover, a permutation test showed that the prediction accuracy in the LT group was significantly lower than in the PLC group (permutation test p=0.02). These results suggest that the fear oddballs induced subjective fear experience and that this reactivity was attenuated following LT.

In addition to the VIFS we applied a neurofunctional signature for unspecific negative affect (PINES). The PINES discriminated fearful vs. neutral oddballs in the LT and PLC group with comparable accuracy (accuracy=79 ±6.2%, binomial test p <0.001, in PLC group; accuracy=77 ±6.3%, binomial test p <0.001, in LT group; permutation p=0.40; **Figure 2D**). These findings, together with the prediction performance of the VIFS, suggest that the fearful oddball can induce a complex array of negative emotional experiences, and LT may have specific effects on subjective fear rather than general non-specific negative affect. Next, we applied the TPS to the data, and the result showed that TPS could not discriminate fearful vs neutral oddballs in either group (PLC: accuracy=47 ±7.6%, binomial test p=0.76 ; LT: accuracy=45 ±7.5%; binomial test p=0.65, **Figure 2E**). These findings were in line with our previous study (23) showing that conditioned fear and subjective fear exhibit distinct neural representations.

We additionally applies these three signatures to classify novel vs. neutral oddballs, and found no group differences (accuracy differences <0.09, Ps >0.26), which further suggest that the effects of LT are fear specific.

### Effects of AT1R blockade on amygdala functional networks

Examination of treatment effects on fearful oddball associated amygdala networks revealed that LT – relative to PLC - significantly reduced bilateral amygdala connectivity with the ventral anterior cingulate gyrus (vACC) during the fearful oddball presentation (peak MNI x, y, z=8, 44, 14; t_85_=-4.30; TFCE p=0.02, FWE-corrected at peak level; k=83, **Fig. 2F**).

## Discussion

Combining pharmacological fMRI with a fearful oddball paradigm and neurofunctional decoding we were able to demonstrate that transient AT1R blockade (1) selectively attenuates neurofunctional reactivity to fear-inducing visual oddballs in terms of attenuating dlPFC activation and amygdala-vACC communication, and (2) specifically reduced subjective fear experience but not the threat or non-specific negative response during fearful oddball presentation. Together the findings indicate that the AT1R system plays a role in neurofunctional fear reactivity and fear experience and that neurofunctional signatures have the potential to improve the precision of pharmaco-fMRI for the characterization of novel receptor targets to alleviate fear.

The conventional mass univariate activation and connectivity analyses revealed that AT1R blockade specifically attenuated left dlPFC reactivity and amygdala-vACC connectivity in response to fearful oddballs. The dlPFC represents a functionally heterogenous region and has been involved in several domains related to fear processing, including implicit and explicit cognitive regulation of negative affect and task-irrelevant distractors (48-56) rapid attention towards salient stimuli, including threat-related stimuli (57-59) and the conscious subjective experience of emotional states including fear (22, 60). In accordance with this functional characterization of the dlPFC in neuroimaging studies lesion and non-invasive brain stimulation studies have provided more causal evidence for an involvement in the fear-related process. Patients with dlPFC lesions exhibited impaired cognitive regulation of subjective fear experience (60) while non-invasive stimulation of this region attenuates vigilance to threatening stimuli (61) and augments fear extinction (62). Likewise the amygdala, the vACC and their interaction have been extensively associated with emotional processing, such that this circuit may encode bottom-up signal indicating the presence of threat and salient information (63) which in turn informs threat acquisition (64) as well as the top-down regulation of emotional distractors (65) and conditioned threat signals (66). Given the functional heterogeneity of the dlPFC and the amygdala-vACC pathway a corresponding interpretation of the specific functional effect of pharmacological ATR1 blockade would remain unspecific. The pattern of a treatment-induced decreased engagement in response to fearful oddballs may for instance reflect decreased automatic threat vigilance, the attenuated experience of fear or a reduced need to engage emotion regulation.

To further determine the specific effects of ATR1 blockade we therefore employed process-specific neurofunctional signatures that have been developed to precisely predict the subjective experience of fear (VIFS), general negative affect (PINES) and conditioned threat reactivity (TPS) (23) based on spatially distributed neurofunctional activation patterns. The VIFS and the PINES successfully discriminated fearful and neutral oddballs in the PLC group, indicating that the fearful oddball induced brain activity patterns reflecting and predictive of subjective fear and general negative affect. Treatment effects were specifically captured by the VIFS such that the subjective fear signature successfully discriminated fearful oddballs from neutral oddballs in the PLC group while ATR1 blockade abolished this discrimination suggesting reduced response of the neurofunctional fear signature. In contrast, the PINES predicting non-specific negative affect successfully discriminate fearful vs. neutral oddballs in both treatment groups, suggesting that while the fearful oddballs per se did not selectively induce fear, ATR1 blockade induced fear-specific effects on the neural level. Notably, the conditioned threat signature did neither discriminate fearful vs. neutral oddballs nor treatment groups confirming a neurofunctional differentiation between conditioned threat and subjective fear experience (see e.g. 22, 23) and arguing against unspecific treatment effects.

Our findings align withevidence suggesting a role of the renin-angiotensin system i.e. the ATR1 in fear and related processes. Rodent models demonstrated that targeting the renin-angiotensin system reduces indices related to fear and anxiety, such that LT-induced ATR1 blockade decreased anxiety as measured by the elevated plus-maze paradigm (67), abolished stress-potentiated (68) and carbon dioxide-induced (41) fear behavior and facilitated threat extinction while it did not affect the acquisition of a conditioned threat response per se (17). Human studies reported sustained amygdala responses to fearful faces as well as increased amygdala regulation via the medial PFC following LT (13, 18) with the present results suggesting a promising potential to decrease the response of the neural signature of subjective fear, that is, reducing the experience of fear.

While the current study provides initial evidence on the potential of LT to attenuate the response of the subjective fear neural signature future studies need to determine its potential in exaggerated fear and to evaluate its potential as an adjunct treatment to reduce and normalize the pathological fear experience.

## Conclusion

Pharmacological blockade of the AT1R attenuated the neurofunctional reactivity to fear-inducing visual oddballs in terms of attenuating dlPFC activation and amygdala-vACC communication, and specifically reduced the response of the subjective fear signature. The findings reflect a functionally relevant role of the AT1R in fear experience which may serve as a promising target for novel interventions to reduce pathological fear.

## Acknowledgements and disclosures

The study was supported by the Brain Project 2022ZD0208500 (MOST2030) of the Ministry of Science and Technology and the National Key Research and Development Program (2018YFA0701400) of China. The authors declare no competing interests.

## Author contributions

FZ and BB designed the study; RZ and ZQ conducted the experiment and collected the data. RZ, WZ, TX, and FZ performed the data analysis. RZ, FZ, and BB wrote the manuscript draft. All authors reviewed and edited the paper.

