## supplements for "Angiotensin II regulates the neural expression of subjective fear in humans - precision pharmaco-neuroimaging approach"

**Zhang et al.,**

**Supplementary material**

**Contact:**

**MRI acquisition and preprocessing**

3.0-T GE Discovery MR750 system (General Electric Medical System, Milwaukee, WI, USA). Structural images were acquired by using a T1-weighted MPRAGE sequence (TR=6ms; TE=2ms, flip angle=9°, field of view=256 × 256 mm2; matrix size=256 × 256; voxel size = 1 × 1 × 1 mm). Functional images were acquired using a T2*-weighted echo-planar sequence (TR=2000 ms; TE=30 ms; slices=39; flip angle=90°; field of view=240 × 240 mm; matrix size=64 × 64, voxel size: 3.75 × 3.75 × 4 mm).

All images were preprocessed and analyzed using Statistical Parametric Mapping (SPM12 v7487, https://www.fil.ion.ucl.ac.uk/spm/software/spm12/ and (1)). The first 5 volumes of each run were discarded to allow for T1 equilibration. In line with our previous studies (1) the remaining volumes were corrected for differences in the acquisition timing of each slice and spatially realigned to the first volume, and unwarped to correct for nonlinear distortions related to head motion or magnetic field inhomogeneity. The anatomical image was segmented into grey matter, white matter, cerebrospinal fluid, bone, fat, and air by registering tissue types to tissue probability maps. Next, the skull-stripped and bias-corrected structural image was generated and the functional images were co-registered to this image. The functional images were subsequently normalized in the Montreal Neurological Institute (MNI) space (interpolated to 2 × 2 × 2 mm voxel size) by applying the forward deformation parameters that were obtained from the segmentation procedure, and spatially smoothed using an 8 mm full-width at half maximum (FWHM) Gaussian kernel.

The first-level general linear model (GLM) included five task regressors corresponding to the 'standard', 'fearful oddball', 'target oddball', 'neutral oddball', and 'novel oddball'. For those participants who missed the target and/or pressed the button during any other conditions, we included additional task regressors to model missing and/or false alarm trials. The fixation-cross epoch served as an implicit baseline. All task regressors were convolved with the canonical hemodynamic response function and a high-pass filter of 128 seconds was applied to remove low-frequency drifts. Motion correction parameters estimated from the realignment procedure were entered as covariates of no interest.

Table 1

| **Stimuli** | **Arousal** | **Valence** | **Fear** |
|---|---|---|---|
| **Standard** | 4.75 (1.80) | 5.96 (1.78) | 2.35 (1.90) |
| **Fearful oddball** | 7.06 (1.52) | 4.76 (2.05) | 3.05 (1.35) |
| **Target oddball** | 4.73 (1.89) | 6.71 (1.42) | 1.57 (0.72) |
| **Novel oddball (30 stimuli)** | 4.48 (0.32) | 6.83 (0.71) | 1.83 (0.61) |
| **Neutral oddball** | 4.18 (0.93) | 6.02 (1.84) | 1.92(1.17) |

*Arousal and valence ratings from original NAPS dataset. Fear ratings from n = 60 independent subjects.

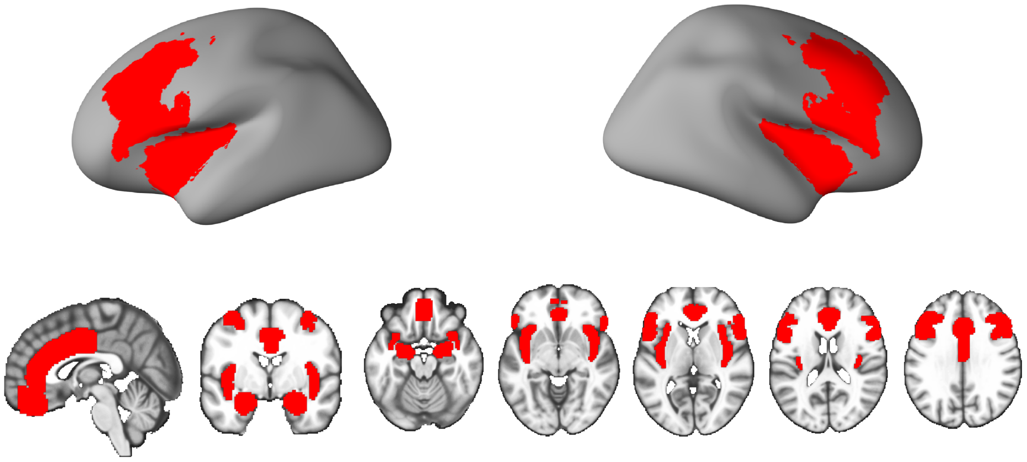

Fig. 1 the mask used for small volume correction (SVC). The mask encompasses the anterior cingulate cortex (ACC), amygdala (Amyg), insula, middle frontal gyrus (referred to as dorsolateral prefrontal cortex (dlPFC)), inferior frontal gyrus (referred to as ventrolateral prefrontal cortex (vlPFC)) and frontal medial cortex (referred to as ventromedial prefrontal cortex (vmPFC)).

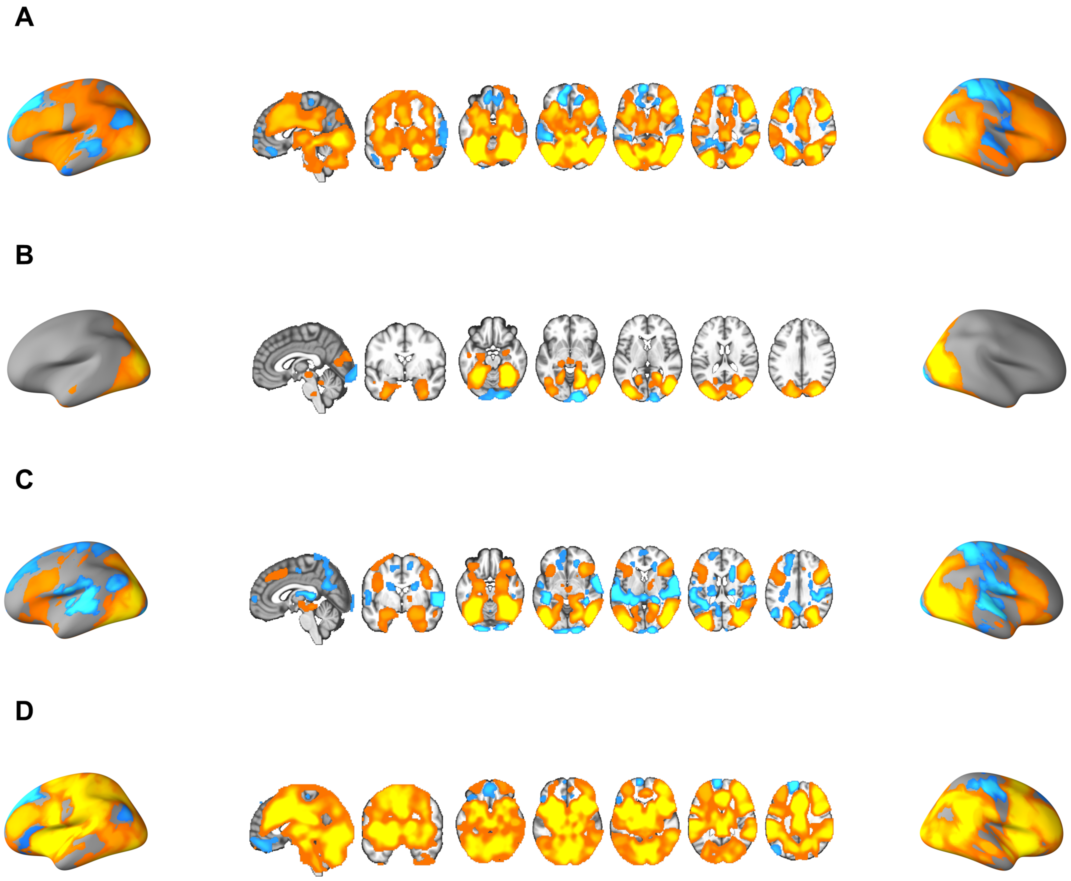


Fig.2 Results of one-sample t test in the PLC and LT group (A) Shows a one sample t-test for the main effect of fearful oddball condition (TFCE $p < 0.05$, FWE corrected); (B) Shows a one sample t-test shows the main effect of neutral oddball condition (TFCE $p < 0.05$, FWE corrected); (C) Shows one sample t-test shows the main effect of novel oddball condition (TFCE $p < 0.05$, FWE corrected); (D) Shows one sample t-test shows the main effect of target oddball condition (TFCE $p < 0.05$, FWE corrected).

1. Zhou F, Zhao W, Qi Z, Geng Y, Yao S, Kendrick KM, et al. A distributed fMRI-based signature for the subjective experience of fear. Nature communications. 2021; 12(1): 1-16.
